# Biodegradable Plastics in the Marine Environment: Comparative Analyses of Bacterial Biofilms, Macroinvertebrate Colonization, and Degradation Patterns

**DOI:** 10.64898/2025.12.22.696091

**Authors:** Kirsten M. Montpetit, Brian T. Nedved, Michael G. Hadfield, Marnie L. Freckelton

**Author notes:** **Corresponding Author:** Marnie L. Freckelton.

## Abstract

Floating debris in the ocean recruits microbes and marine invertebrates to its surface, resulting in rafting communities. As biodegradable plastics increase in prevalence as alternatives to petroleum-based plastics, their properties may impact the dynamics of rafting communities by influencing the composition of bacterial biofilms and attached invertebrates. We compared attached biofouling communities of bacteria and marine invertebrates on biodegradable surfaces and nonbiodegradable petroleum-based plastics and naturally occurring substrata. Six surface types: polypropylene plastic (PP), polystyrene (PS), polylactic acid plastic (PLA), polyhydroxyalkanoates plastic (PHA), maple wood veneer, and propagules of the mangrove *Rhizophora mangle*, were examined to determine if plastic type affected biofilm composition on the surfaces and their degradation patterns. Biofilm analyses were conducted at twelve weeks, and degradation analyses were conducted at twenty-two weeks of immersion. Using digital image analysis, 16S rRNA sequencing, and metagenomic analyses, we found that microbial biofilms, marine invertebrate inhabitants, and degradation patterns differed across the various substrate types tested. Microbial communities on PLA were more similar to those on the two non-biodegradable plastics, while communities on PLA were more like those on the natural substrata. The biodegradable PLA showed signs of degradation within 22 weeks of immersion, suggesting that biodegradable plastics behave variably in the natural environment. Results of this study bring forth the importance of designing biodegradable plastics with careful consideration of the environmental conditions in which they are likely to persist.

## 1. Introduction

Marine rafting communities are diverse assemblages that develop on floating substrates and facilitate the long-distance dispersal of native and non-native taxa through passive transport (Thiel & Gutow, 2005). While such rafting enhances biogeographic connectivity, it also increases the risk of biological invasions (Goldstein et al., 2014; Kiessling et al., 2015; Rech et al., 2016). Rafting communities also influence nutrient cycling, predator-prey interactions, and biodiversity (Goldstein et al., 2014; Kiessling et al., 2015; Rech et al., 2016; Thiel & Gutow, 2005). These communities colonize natural substrates, such as seeds, pumice, and driftwood, as well as anthropogenic debris, including plastic waste, fishing gear, and vessel components (Kiessling et al., 2015; Rech et al., 2016). Understanding how rafting communities assemble and change is crucial for predicting ecological impacts and informing conservation and biosecurity efforts.

The colonization process of floating substrates is similar to that on fixed benthic habitats (Pinochet et al., 2019). First, microorganisms rapidly colonize and form diverse biofilms (Hadfield et al., 2025; Zobell & Allen, 1935). These biofilms then produce cues to trigger attachment and metamorphosis of marine invertebrates (Cavalcanti et al., 2020; Dobretsov & Rittschof, 2020; Hadfield, 2011; Hadfield et al., 2025; Hadfield & Paul, 2001). However, biofilm communities vary in composition depending on substrate type, exposure time, and environmental conditions (Lazcano et al., 2024). These factors shape chemical landscapes that can either promote or inhibit larval attachment (Dobretsov & Rittschof, 2020; Pinochet et al., 2019). As a result, the type of floating substrate can affect not only biofilm composition but also the identity, diversity, and timing of invertebrate colonization.

Due to inadequate and improper disposal methods, floating substrates in the world’s oceans are now dominated by anthropogenic materials, especially plastic debris (Bergmann et al., 2015; Pinochet et al., 2019; Walker, 2017). Each year, an estimated 8-12 million metric tons of plastic enter marine environments (Jiang et al., 2018). The World Wide Fund for Nature reported in 2023 that 70 percent of marine plastic debris entering the ocean consists of single-use containers. These containers, such as bottles and to-go boxes, can trap air and float on the ocean’s surface even if they are denser than seawater. Common petroleum-based plastics, such as polypropylene (PP) and polystyrene (PS), are non-biodegradable and, once introduced into the marine environment, accumulate and will remain unless physically removed (Geyer et al., 2017; Tokiwa et al., 2009). These plastics will gradually fragment into micro- and nanosized particles under continued exposure to UV radiation, salinity, temperature, physical abrasion, and microbial activity (Barnes et al., 2009; Dimassi et al., 2022; Lv et al., 2024). The rate and extent of fragmentation are further influenced by polymer chemistry, molecular weight, and hydrophobicity (Dimassi et al., 2022). This fragmentation contrasts sharply with many biological substrates that degrade on far shorter timescales; e.g., a wooden log is estimated to degrade within 2 years (Jiang et al., 2018).

Humans incorporate plastic into much of their daily routines, relying on it so heavily that, despite widespread awareness of its severe environmental consequences, global plastic production and consumption continue to increase (Pilapitiya & Ratnayake, 2024). Single-use plastics are widely adopted for convenience, yet their disposal pathways are often inconvenient. Bioplastics are plastics derived from renewable biological sources, also referred to as bio-based. Bioplastics are increasingly marketed as alternatives to petroleum-based plastics in an effort to reduce the negative ecological impacts of the ever-growing demand for single-use disposable plastics (Di Bartolo et al., 2021; Tokiwa et al., 2009). Many bioplastics are also biodegradable, meaning they break down through biological processes. While the definitions of biodegradable and bioplastic are distinct, they are often used interchangeably. Here, we examine the colonization and degradation of two biodegradable bioplastics, polylactic acid (PLA) and polyhydroxyalkanoates (PHA). PLA is derived from sugar and starch, whereas PHA is produced from bacterial by-products; PLA and PHA are among the major types of biodegradable bioplastics currently being produced (Acharjee et al., 2022; Lackner, 2015; Naser et al., 2021). PLA and PHA are used as alternatives to petroleum-based plastic polymers in packaging materials and other applications aiming to reduce environmental impact (Atarés et al., 2024; Auras et al., 2004). The use of both PLA and PHA is projected to continue to increase in the coming years (Ali et al., 2023; European Bioplastics, 2022; Shi et al., 2025).

Despite their promise as alternatives to conventional plastics, there is limited knowledge on the fate of biodegradable plastics in the marine environment. Current international and American standards and certifications (ASTM, 2019; ISO, 2005; ISO, 2016; ISO, 2018; ISO, 2019) used to determine biodegradable status rely on standardized laboratory tests conducted under controlled conditions. While useful for benchmarking materials, these tests may not accurately reflect the complexity and variability of natural marine environments (Brožová et al., 2025). Multiple pathways for degradation exist, with the rate dependent on chemical and physical properties, environmental conditions, and microbial activity (Tokiwa et al., 2009). This limitation is particularly relevant for PLA, whose hydrolysis-driven, temperature-dependent degradation is not directly measured by current ISO and ASTM-standardized tests that primarily measure biodegradation by carbon dioxide breakdown (ASTM, 2019; Huang et al., 2025; ISO, 2005; Limsukon et al., 2023; Ranakoti et al., 2022). As a result, current certifications may not accurately represent how biodegradable plastics behave in natural marine settings. Given both the projected increase in use and the inconsistencies in the behaviors and properties of biodegradable materials, it is important to investigate the marine microbes and invertebrates colonizing biodegradable plastics and the degradation of these materials in the ocean.

Hawai‘i is a particularly vulnerable victim of marine plastic pollution due to its proximity to gyres and convergence zones in the North Pacific, where significant accumulation of garbage occurs, such as the Great Pacific Garbage Patch (Howell et al., 2012). A study by Lebreton and colleagues (2018) reported that plastic debris in the Great Pacific Garbage Patch is increasing exponentially compared to surrounding waters. As a result, the majority of marine plastic debris found in Hawai‘i has floated there from distant regions (Brignac et al., 2019). To combat the detrimental environmental impacts of plastic pollution, Hawai‘i has banned single-use plastics, including polystyrene foam containers and plastic bags. Instead, people are encouraged to switch to biodegradable materials (City and County of Honolulu, 2019). Although these plastics should be disposed of at proper facilities, when in the hands of the general public, they often enter the natural environment. It can reasonably be assumed that these biodegradable plastics will increasingly be found in our oceans as they grow in popularity.

The research presented here examines how substrate type influences the dynamics of rafting communities, including microbial colonization and macroinvertebrate settlement. The study directly compares non-biodegradable petroleum-based plastics, biodegradable bioplastics, and naturally occurring materials commonly found in marine environments (propagules of *Rhizophora mangle* and maple wood veneer). Three questions guided the investigation: (1) Do plastic polymer types differ in their degradation patterns over time? (2) Do different substrates develop distinct early- and late-stage bacterial biofilm communities? and (3) Does variation in bacterial biofilm composition drive differences in the colonization and development of early-stage marine invertebrate communities? This comparison provides insight into the potential of marine biofilms to influence rafting communities across substrate types. A comprehensive understanding of how biodegradable plastics impact these marine communities will aid in the development of effective strategies to mitigate and manage marine plastic pollution as biodegradable plastics become increasingly abundant.

## 2. Materials and Methods

### 2.1 Experimental Fouling Surfaces

Six substrate types were compared: four plastics — polypropylene (PP), polystyrene (PS), polylactic acid (PLA), and polyhydroxyalkanoate (PHA) — and two biologically derived materials, maple veneer (WOOD) and mangrove propagules of *Rhizophora mangle* (PROP). The non-biodegradable petroleum-based plastics (PP and PS) were cut from transparent food-storage containers (Promoze; SafePro). PP sheets (88 × 130 × 0.25 mm) and PS sheets (97 × 97 × 0.25 mm) were prepared from the largest uniformly flat sections available from each container type, ensuring consistent dimensions across replicates while avoiding warped or molded areas. Polymer type was verified based on the manufacturer’s specification for each specific product, as well as by secondary verification at Ho□omalu Ke Kai’s plastic upcycling facility (Kaua□i, Hawai‘i) using a plastic verification device (PlasTell Desktop; Matoha, U.K.). The biodegradable bio-based substrates PLA (Overture Plastics, Texas, USA) and PHA (Beyond Plastic, California, USA) were produced by 3D printing (Prusa i3 MK+ printer) using white filament. Commercial products marketed as PLA or PHA frequently contain undocumented polymer blends or additives; therefore, printing ensured material composition and reproducibility. Printed panels (110 × 159 × 0.25 mm) represented the largest dimensions that could be reliably fabricated without warping or surface defects. As a result of the printing process, the PLA and PHA substrates were slightly textured. The maple veneer panels (110 × 159 × 0.25 mm; Sauers & Company Veneers) matched the dimensions of the PLA and PHA panels. Veneer panel size was determined by the maximum uniform area obtainable from the vendor-supplied sheets while avoiding knots or grain deformities that could affect surface structure. The mangrove propagules of *Rhizophora mangle* (approximately 14 mm diameter × 130 mm length) were collected from living trees along the water’s edge at Neal S. Blaisdell Park in Pearl Harbor (21.38455° N, 157.95359° W). Only propagules hanging above the water were collected to ensure they had not been previously submerged and fouled. Propagules were included because they represent a naturally occurring component of the floating debris community in Hawaiian coastal waters and provide an ecologically relevant comparison for colonization dynamics. Their dimensions were not modified, as preserving their natural size and shape was essential to maintaining the integrity of this biological substrate. While substrate source and usable area varied among substrates, dimensions were standardized within each material, and all analyses were normalized to surface area to enable direct comparison of biofilm development.

### 2.2 Field Site

The test surfaces were submerged in water from a pier on Ford Island in Pearl Harbor (21.35745°N, 157.96018°W) (Figure 1A). Pearl Harbor is an active warm-water U.S. Navy port (temperatures during this study ranged between 24°C and 27°C) with year-round biofouling, allowing the larvae of biofoulers to settle on submerged surfaces (Holm et al., 2008).

**Figure 1.**
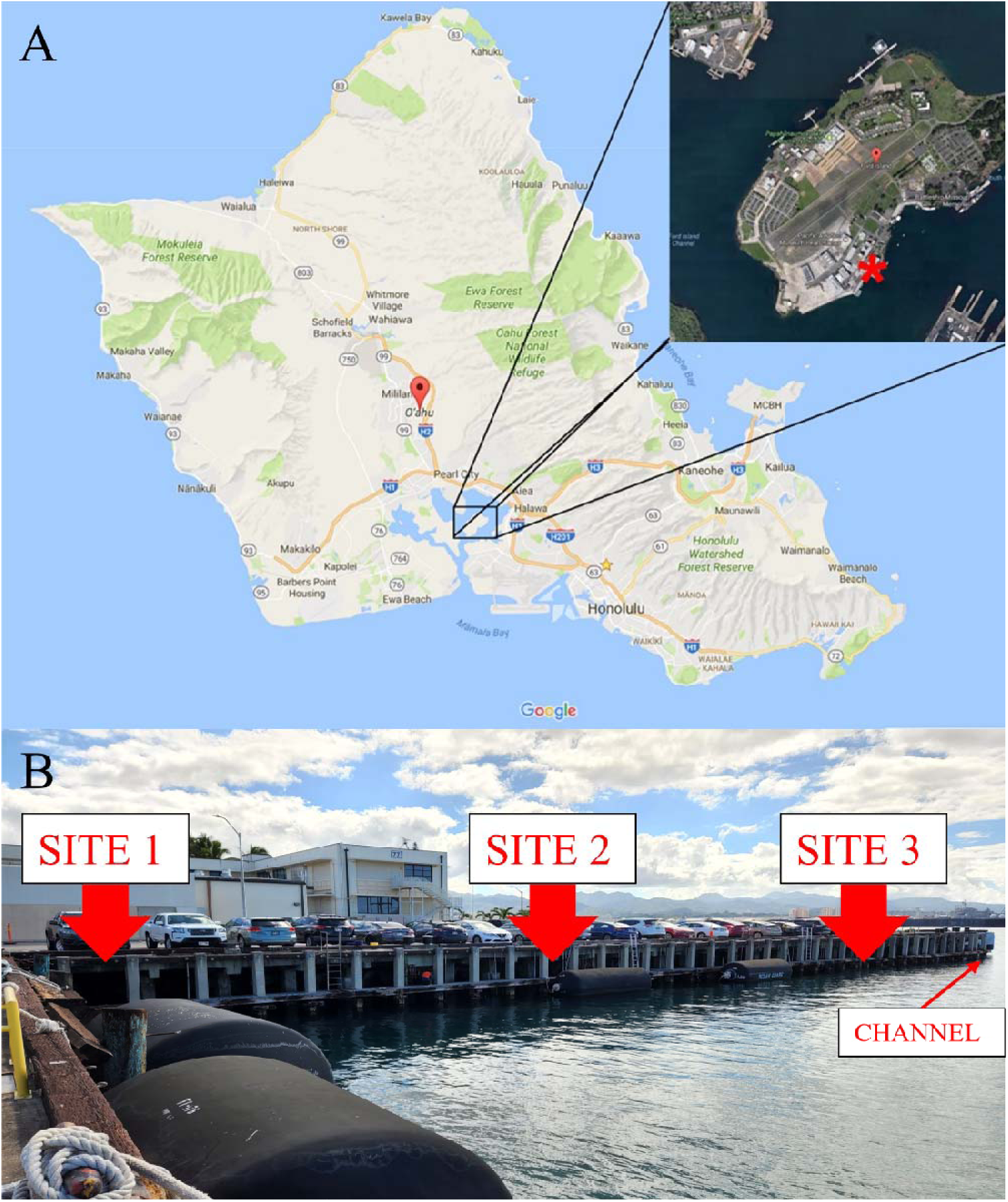
Map and Study Sites at Pearl Harbor, Oϑahu. (A) A map of O‘ahu, Hawai‘i, showing the location of the study site (red star) on Ford Island within Pearl Harbor, Hawai‘i. (B) The pier from which the experimental surfaces were submerged in Pearl Harbor. Red arrows indicated the sites where the cages containing the substrates were suspended along the pier (Site 1, Site 2, Site 3), adjacent to the harbor boat channel and pier.

### 2.3 Experimental Design

To study the settlement and development of early-stage marine biofouling communities on plastics in the field, three cages were built, each 1.3 meters long, 0.4 meters wide, and 0.4 meters tall. Each cage consisted of a PVC pipe frame covered with Vexar mesh (6.35mm by 6.35mm) to prevent fish from grazing on the experimental surfaces (Figure 2). Substrates were randomly assigned positions within the cages, with one of each substratum type per cage (Figure 2). Cages were suspended approximately 1 meter below the mean low tide and deployed along the length of the dock, at three different distances from the ship channel (Figure 1B). This experiment was conducted in two phases spanning over 22 weeks, from October 17, 2023, to March 21, 2024. The first phase included microbial and macrofouling analyses conducted over twelve weeks. Photos of each surface were taken weekly for the first four weeks and then biweekly for the remaining eight weeks for macrofouling analysis. Bacterial samples were collected at week one and week twelve of the experiment. Upon completion and collection of results of the first phase, we elected to continue the degradation experimentation for another ten weeks to maximize exposure time, allowing for longer-term analysis. Due to logistical constraints, biofouling analyses were not conducted again at 22 weeks.

**Figure 2.**
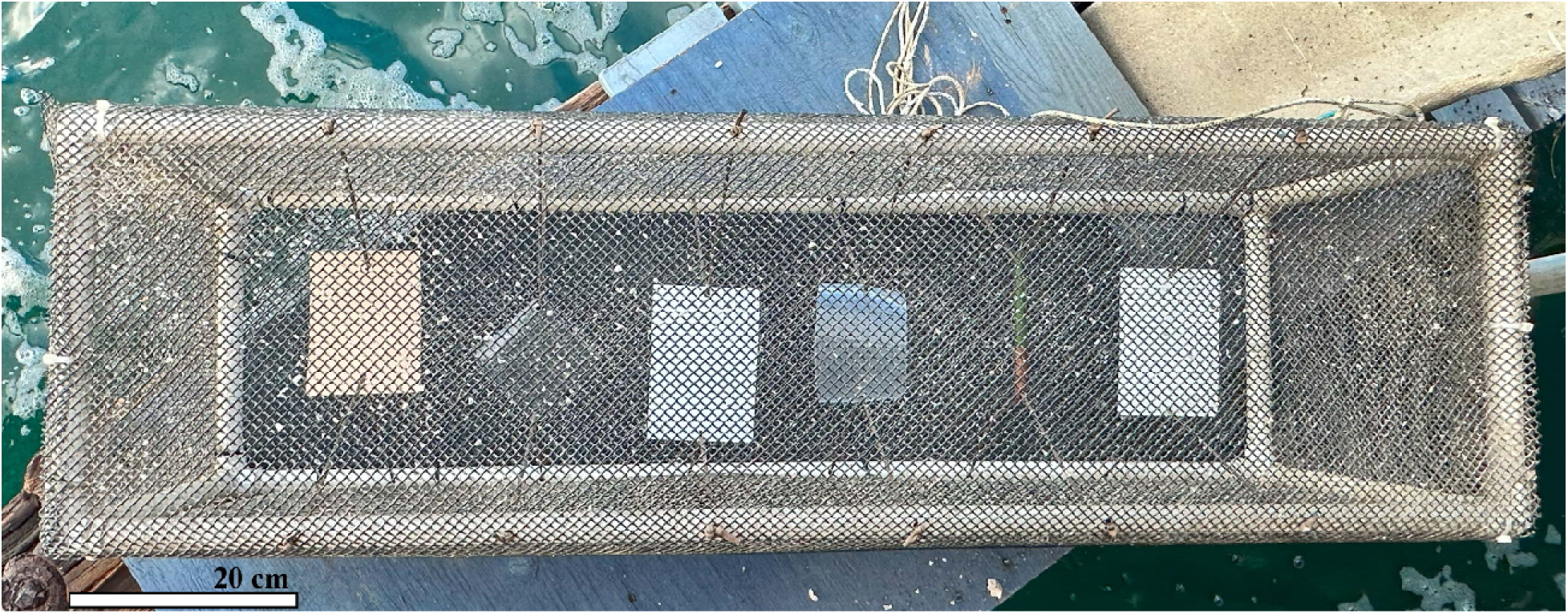
Cage with Suspended Substrates. One of the constructed cages with the six substrate types suspended within it.

### 2.4 Analysis of Substrate Degradation

The force required to penetrate each substratum was measured using a chisel tip (0.28 mm x 8.91 mm) on a SHIMPO Mechanical Force Gauge with a maximum capacity of 20 pound-force (lbf). The value was then converted to SI units (pascals) for analysis. Each substratum was clamped over a wooden block with a milled-out hole that was larger than the chisel tip, as shown in Figure S1 (See supplementary material). Penetration was measured at three, evenly spaced points per replicate and averaged. For the wood veneer, measurements were recorded with the chisel tip aligned with and against the grain of the veneer. Penetration forces were measured for each material before exposure to seawater and on those that had been immersed for 22 weeks. Three replicates of each material were assessed and averaged for the pre-immersion measurements and post-immersion measurements when possible.

### 2.5 Biofilm Sampling, DNA Extraction, and 16S rRNA Sequencing

Microbial biofilm communities were sampled from each substratum after one week and twelve weeks of immersion. At each time point, biofilms were collected from all substrate replicates using dry sterile swabs, as they are non-destructive to the substrate and communities present. Samples were collected via swab using a standardized side-to-side motion along a consistent area down the surface of each substrate. Swabs were immediately fixed in 95% ethanol and stored at –80°C until processing. DNA was extracted using the Qiagen PowerSoil Kit (Venlo, Netherlands) with minor modifications for improved lysis and yield. Swabs were transferred into the bead-containing lysis tubes supplied with the kit, briefly vortexed, and incubated at 37°C for 20 minutes to enhance cell disruption. Samples were then homogenized using a bead mill (Fisherbrand Bead Mill 24 Homogenizer) for 30 seconds, paused for 30 seconds, and homogenized for an additional 30 seconds. Finally, after the standard binding and washing steps, the silica membrane in the spin column was dried briefly in an oven at 37°C to remove residual ethanol, and the DNA was eluted with DNase-free water (2 x 20 µL). Extracted DNA was stored at –80°C. DNA concentration and purity were assessed using a NanoDrop spectrophotometer (Thermo Scientific NanoDrop 2000). 16S rRNA gene libraries were prepared from 10 ng of template DNA using the Oxford Nanopore 16S Barcoding Kit (SQK-RBK114.24). The MinION PCR protocol was followed with an increased cycle number of 30 rather than the recommended 25. Early-stage marine biofilms yielded low DNA quantities, and preliminary tests showed that 25 cycles did not reliably produce sufficient amplicons for library construction. Increasing to 30 cycles improved amplification success while remaining within a range that minimized amplification bias and chimera formation. Libraries were sequenced on a MinION Mk1B (Oxford Nanopore Technologies, UK). Bacterial community composition was characterized using the preconfigured 16S rRNA workflows (v1.5.0) in EPI2ME (Oxford Nanopore, v5.2.3). Reads with <80% sequence similarity and taxa with fewer than five reads per sample were excluded to improve taxonomic reliability and reduce noise from poorly supported assignments in accordance with standard practice.

### 2.6 Composition Analysis of Macrofouling Community

Photos of the experimental panels were taken weekly from the same relative positions. The composition of macrofouling communities that colonized each surface was analyzed using digital imaging and computer image analysis (Photoquad, Version 1.4, Trygonis and Sini, 2012). Differences in percent coverage by marine invertebrates were compared across substrata by overlaying fifty points onto the images using a stratified random design. The types of fouling organisms (See supplementary material, Table S1) on the surfaces of the panels were estimated for each point. If no foulers were present under the point, but diatoms had settled onto the surface or sediment build-up was observed, it was assigned to the slime category. If neither diatom films nor macrofoulers were present beneath the point, it was assigned to the bare category.

### 2.7 Statistical Data and Diversity Analyses

Upon obtaining all data from microbial and macrofouling analyses, non-metric multidimensional scaling (NMDS), Shannon diversity, and permutational multivariate analysis of variance (PERMANOVA run via Adonis) analyses were run on each dataset using statistical packages Vegan (Oksanen et al., 2022), MASS (Venables & Ripley, 2002), and tidyverse (Wickham et al., 2019) in RStudio (Posit Team, 2024; R Core Team, 2024). The Bray-Curtis dissimilarity statistic was used to visualize the variation in community composition among substrate groups and to generate the NMDS for each data set. A PERMANOVA analysis based on distance matrices was chosen to determine the statistical significance of the groupings and the differences in overall community structure for both the macrofouling and microbial analyses.

### 2.8 Data Availability

All data in this manuscript are deposited in Figshare and are publicly available. https://figshare.com/s/def15910cb8aa5e5e68b

## 3. Results

### 3.1 Substrate Degradation

In the pre-immersion plot (Figure 3A), it showed that the forces required to puncture PP, PLA, and the propagule were near or at the maximum force measurable by the force gauge (37.5 MPa). In the post-immersion plot (Figure 3B), there was a minimal reduction in the mean force required to penetrate the PP panel, decreasing by only 0.40 MPa. The PS panel also showed only a slight decrease of 3.2 MPa in the mean penetration force. In contrast, by week 22, the propagule became increasingly fragile over time, with a 27 MPa decrease in mean force to penetrate, and it displayed visual signs of breakdown. The non-biodegradable plastics, PP and PS, did not visibly appear to degrade. The wood displayed a similar pattern with a 16 MPa decrease when in line with the grain. The substrates, wood and mangrove propagule, that became too fragile to determine force-gauge values were excluded from the post-immersion plot. One biodegradable plastic, PLA, showed a significant reduction in the mean force required to penetrate over time, decreasing by 1.0 MPa (p<0.05; See supplementary material, Table S2). PHA displayed a unique pattern, requiring more force (21 MPa at week one and 25 MPa at week 22) to penetrate after 22 weeks of immersion than it did at one week of immersion, but this difference was not significant (Figure 3).

**Figure 3.**
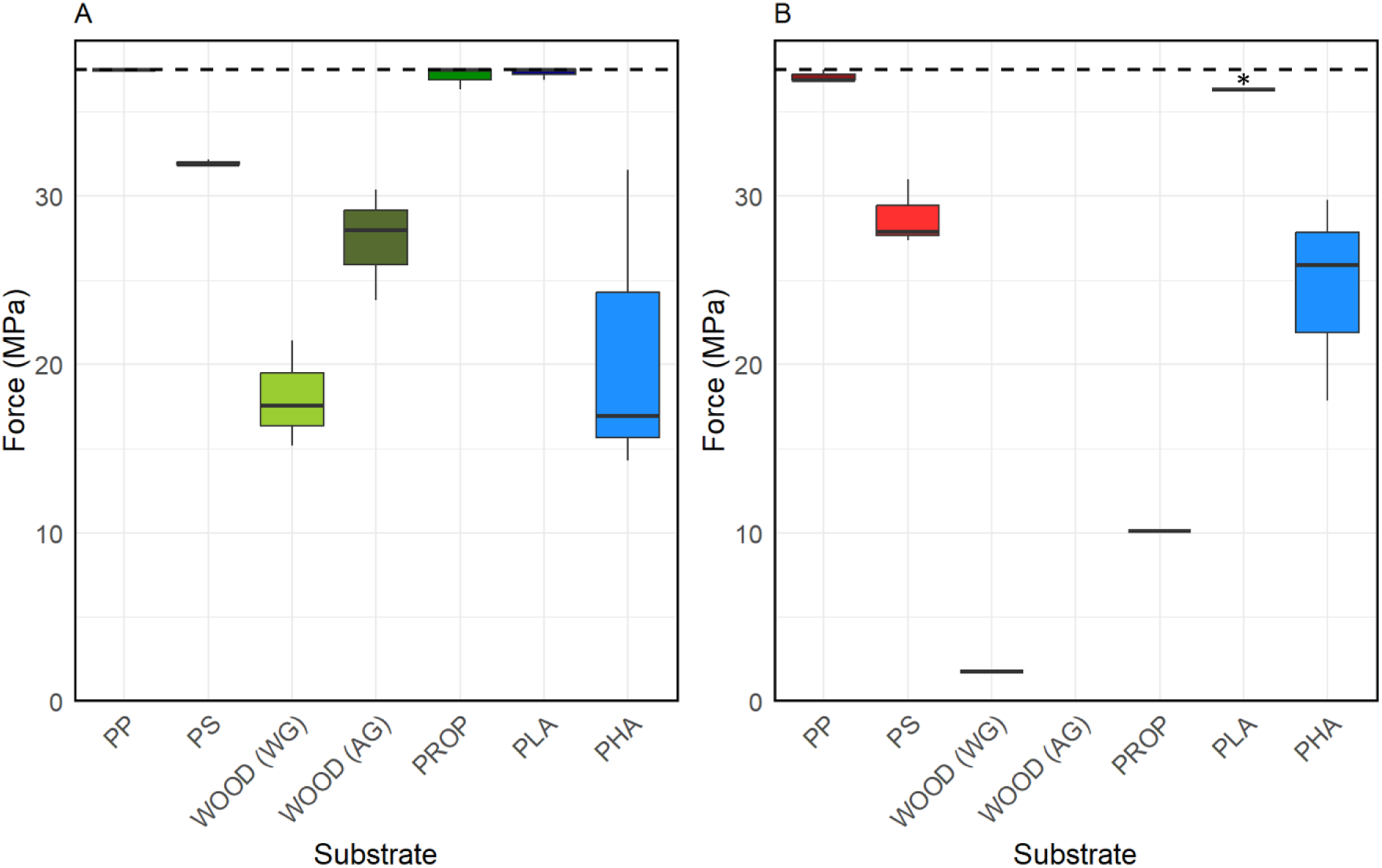
Measurements of Penetration Force through each Substrate. The force required to penetrate each experimental substrate before (A) and after (B) 22 weeks of immersion in Pearl Harbor was measured with a force gauge, with force in megapascals (MPa) on the y-axis and substrate type on the x-axis. The dotted line near the top of each plot indicates the maximum value that the force gauge was able to measure. Substrates tested included Polypropylene (PP), Polystyrene (PS), Polylactic Acid (PLA), Polyhydroxyalkanoate (PHA), mangrove propagules from *Rhizophora mangle* (PROP), and wood maple veneer tested with the grain [WOOD (WG)] and against the grain [WOOD (AG)].

### 3.2 Bacterial Diversity

Sequencing of marine biofilm bacterial communities from plastics, wood, and mangrove propagules generated a total of 5,037,797 Amplicon Sequence Variants (ASVs). After rarefaction, 34,190 sequences per sample were analyzed (See supplementary material, Figure S2). The total number of ASVs identified at a 97% DNA similarity threshold was 3,451. Three samples collected in week 12 (PP, PS, PLA) did not meet the rarefaction threshold and were therefore excluded from further analysis. Longer substrate immersion time corresponded with more diverse microbial communities. Alpha diversity of the microbial populations (Shannon diversity index) increased over the twelve weeks for all substrates (F=17.72, p<0.05) (See supplementary material, Table S3). Biological and biodegradable substrates showed greater increases in diversity than PP and PS, with the largest increases observed in the propagule, wood, and PLA biofilms (Figure 4); by week 12, the wood biofilms had the highest overall alpha diversity of all the treatments (Figure 4; See also supplementary material, Table S3). Traditional plastics, PP and PS, had a less substantial increase in alpha diversity from week one to week twelve compared to the increase observed for biological and biodegradable substrates (Figure 4). Analysis of the ten most prominent bacterial genera present on each substrate type yielded a total of eighteen bacterial genera (Figure 5). Eleven of the eighteen genera are from the phylum Cyanobacteriota, followed by three in the phylum Campylobacterota, and one each in the phylum Pseudomonadota, Bacillota, Thermodesulfobacteriota, and Myxococcota (Figure 5). The genera *Prochlorococcus* and *Fisherella*, both in the phylum Cyanobacteriota, consistently dominated biofilms on all substrates at both time points (Figure 5). *Leptochromothrix*, also belonging to the phylum Cyanobacteriota, appeared in high abundance across all substrates in week one but declined by week twelve (Figure 5).

**Figure 4.**
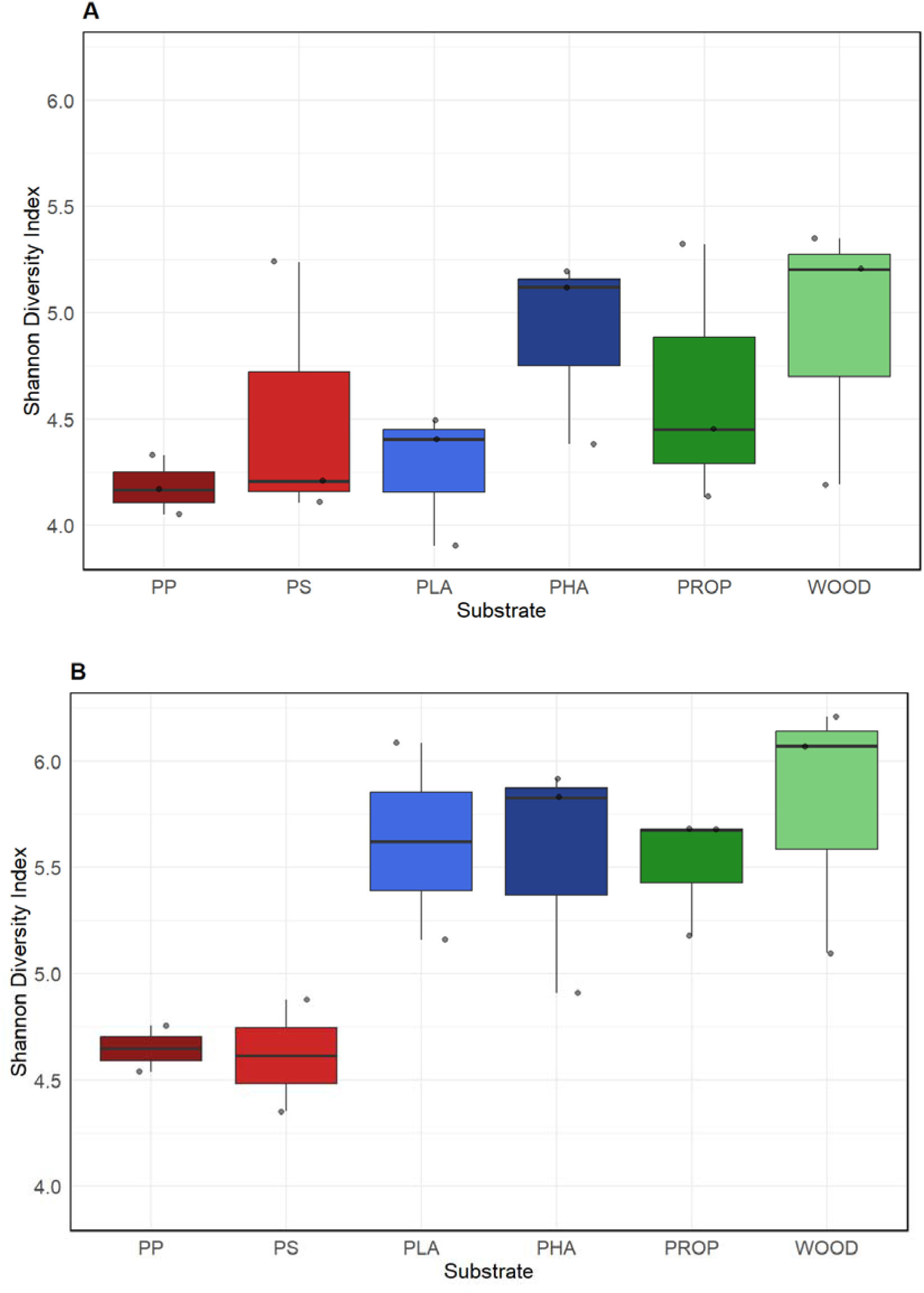
Shannon Diversity Indices for Microbial Communities. Species richness and evenness were measured using Shannon Diversity Index (SDI) analyses for each substrate type at week one (A) and week 12 (B). A larger SDI value corresponds to higher alpha diversity, while a smaller value indicates fewer dominant species and lower alpha diversity. Substrate types include non-biodegradable plastic polymers, Polypropylene (PP) and Polystyrene (PS), biodegradable plastic polymers, Polylactic Acid (PLA) and Polyhydroxyalkanoate (PHA), and biological materials, mangrove propagules from *Rhizophora mangle* (PROP) and wood in the form of maple veneer (WOOD).

**Figure 5.**
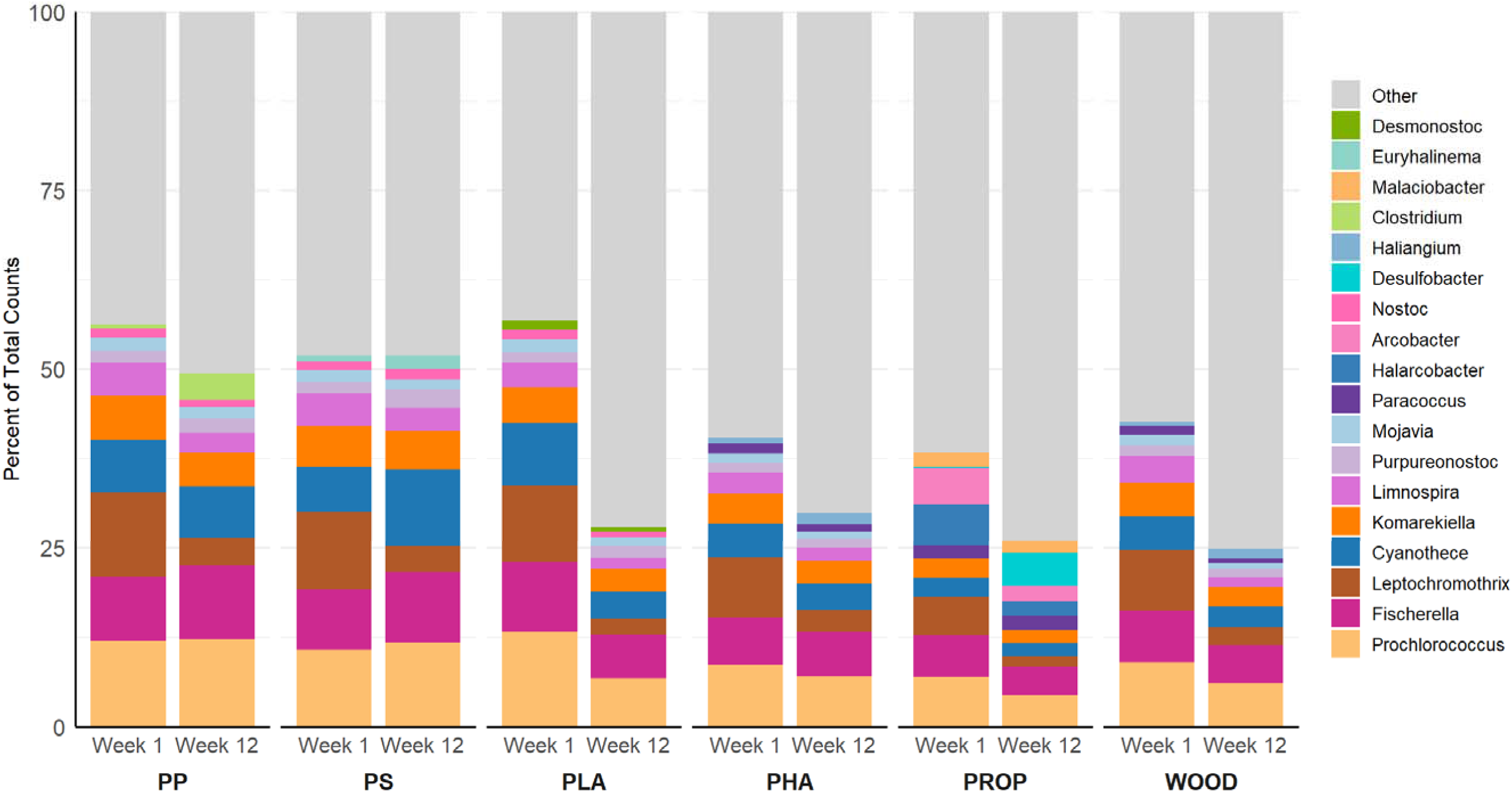
Relative Abundance of the Top Ten Most Prevalent Bacterial Genera on Each Substrate. A stacked bar plot visualizing the top ten most prevalent bacterial genera on each substrate type at week one and week twelve; all remaining low-abundance ASVs are included in the “other” category. Substrate type and time are shown on the x-axis and relative abundance proportions on the y-axis, with colors corresponding to various bacterial genera. Substrates include Polypropylene (PP), Polystyrene (PS), Polylactic Acid (PLA), Polyhydroxyalkanoate (PHA), mangrove propagules from *Rhizophora mangle* (PROP), and wood maple veneer (WOOD).

Over time, more rare taxa increased in relative abundance, contributing to the increased “other” category observed in week twelve (Figure 5). *Clostridium* (phylum Bacillota) appeared exclusively on PP, increasing from 0.5% in week one to 3.7% by week twelve (Figure 5). *Euryhalinema* (phylum Cyanobacteriota) was observed only on PS with a slight increase in relative abundance from 0.8% to 1.9% in the twelve weeks of exposure (Figure 5). Despite these distinct generic differences, PP and PS supported similar overall microbial communities (Figure 6). On PLA, the proportion of the top ten abundant bacteria declined between week one and week twelve, corresponding to an increase in the proportion of rare ASVs (Figure 5). PLA also recruited *Desmonostoc* (phylum Cyanobacteriota), which was absent from all other substrates (Figure 5). *Nostoc* was consistently detected among the top ten taxa on PP, PS, and PLA, but was absent from PHA, the propagule, and wood microbial communities (Figure 5). In contrast, *Paracoccus*, the only member of the Pseudomonadota among the dominant genera, appeared on PHA, the propagule, and wood, but not on PP, PS, or PLA (Figure 5). *Haliangium*, the only member of the phylum Myxococcota, was found in low abundance on both PHA (0.86% in week one, 1.6% in week 12) and wood (0.56% in week one, 1.4% in week 12) (Figures 5 & 6). The propagule supported a more diverse microbial community, colonized by multiple unique taxa, including *Malaciobacter* (phylum Campylobacterota)*, Arcobacter* (phylum Campylobacterota), and *Desulfobacter* (phylum Thermodesulfobacteriota). *Desulfobacter* increased substantially from 0.2% in week one to 4.7% in week twelve (Figure 5). Finally, PHA and wood shared the same top ten ASVs in comparable abundances (Figure 5).

**Figure 6.**
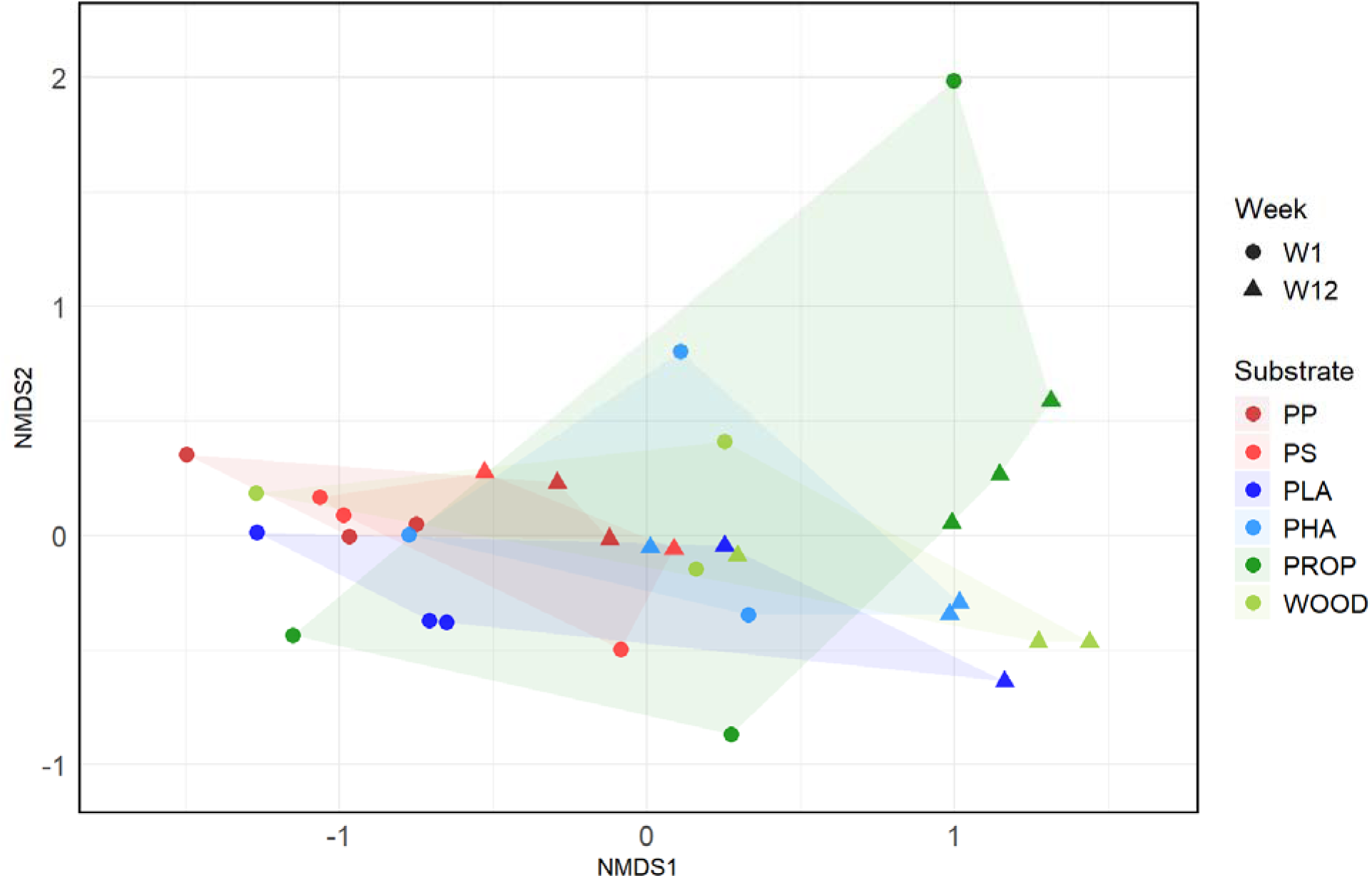
NMDS Bray-Curtis Ordinance Plot for Microbial Communities. Non-metric multidimensional scaling (NMDS) plot that displays differences in the microbial communities present on the substrate types, indicated by different colors. Substrate types are Polypropylene (PP), Polystyrene (PS), Polylactic Acid (PLA), Polyhydroxyalkanoate (PHA), mangrove propagules from *Rhizophora mangle* (PROP), and wood maple veneer (WOOD). Circles represent week one samples; triangles represent week 12 samples. The more clustered the points are together, the more similar their microbial communities are. Shading shows the extent of variability in the biofilms present relative to the centroids of each substrate.

Non-metric multidimensional scaling analysis (NMDS) based on a Bray-Curtis distance matrix revealed distinct microbial community compositions across substrates (Figure 6). Notably, the microbial community on PLA, a biodegradable plastic, clustered with non-biodegradable plastics, PP, and PS. In contrast, the other biodegradable plastic, PHA, hosted a community more similar to those on the biological materials, the propagule and wood (Figure 6). PERMANOVA analysis comparing substrate type (biodegradable, biological, non-biodegradable) revealed a significant effect on microbial community composition (R² = 0.41, F_27,5_ = 3.82, p < 0.05) (See supplementary material, Table S4). Although the variability between microbial communities on the different substrates was not statistically significant following the multiple testing correction, ordination patterns and raw pairwise PERMANOVA results suggest that PHA shares more in common in terms of microbial community composition with biological substrates, while PLA shares more with the non-biodegradable substrates.

### 3.3 Marine Invertebrate Diversity

Macrofouling communities differed between biological substrates and non-biodegradable plastics (Figure 7). The two biodegradable plastics differed in invertebrate community composition, with PLA communities being more similar to those on non-biodegradable plastics, and PHA supporting communities more similar to those on biological substrates (Figure 7). Immersion time also influenced macrofouling community development (Figure 7). Across all substrates over the twelve weeks, bare surfaces and slime or sediment cover remained the dominant fouling category, and calcareous tubeworms were consistently major colonizers (Figure 8). PHA accumulated a greater number of bivalve molluscs over time, while PLA and PP followed a similar pattern but at lower abundances. The mangrove propagule had colonial and solitary tunicates in earlier weeks and encrusting bryozoans in later weeks (Figure 8). In addition to calcareous tubeworms, molluscs were also present on the PHA and wood (Figure 8). Macrofouling communities on PP, PS, and PLA showed higher taxonomic variability in macrofoulers’ composition than what was seen on PHA, wood, and the mangrove propagule (Figure 8).

**Figure 7.**
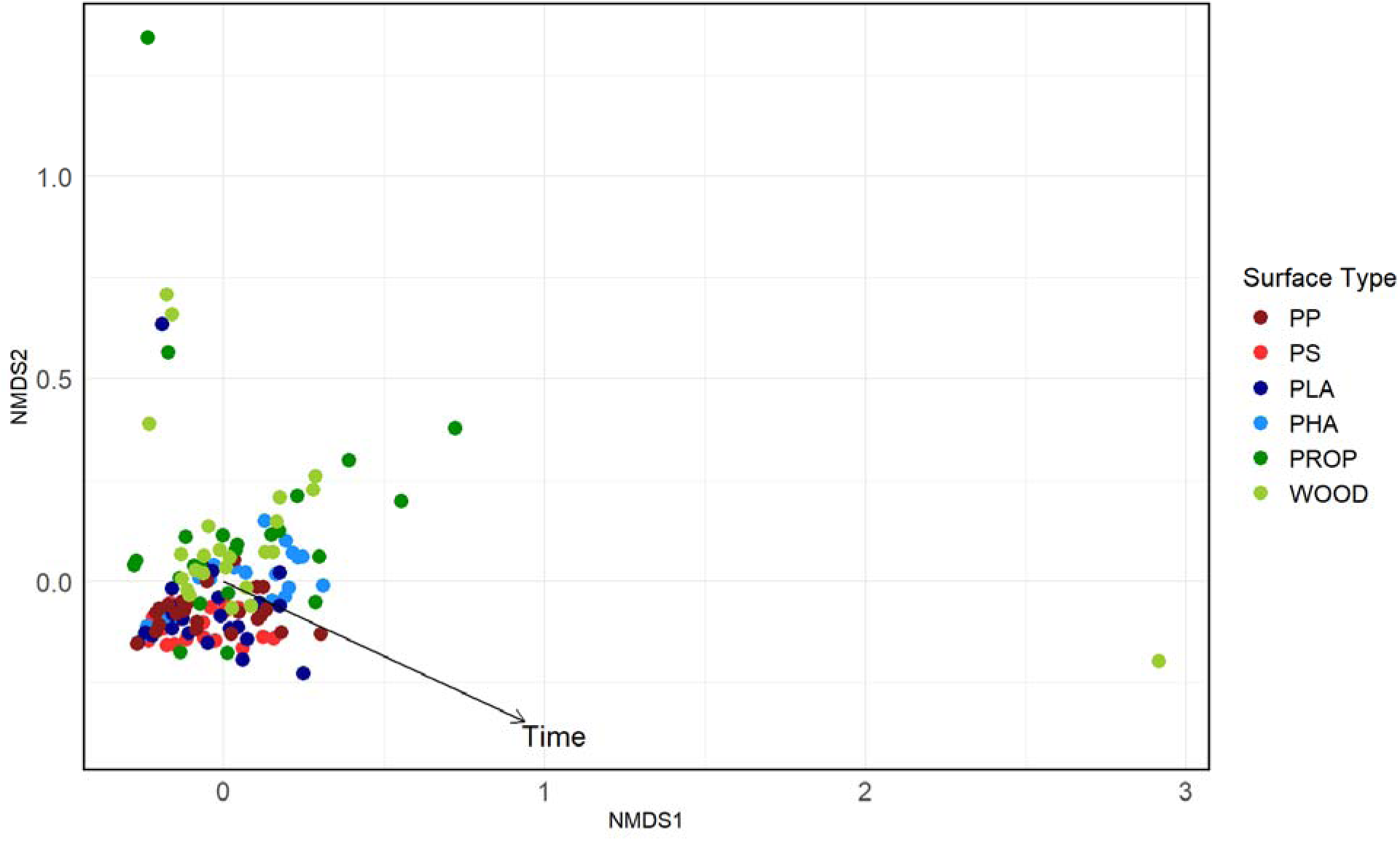
NMDS Bray-Curtis Ordinance Plot for Macrofouling Communities. Non-metric multidimensional scaling (NMDS) plot showing differences in macrofouling communities across substrate types, indicated by different colors. The vector for the effect of time on the macrofouling communities is shown as an arrow. Points closer together are more similar, and the longer the arrow, the stronger the effect. The substrates analyzed include Polypropylene (PP), Polystyrene (PS), Polylactic Acid (PLA), Polyhydroxyalkanoate (PHA), mangrove propagules from *Rhizophora mangle* (PROP), and wood maple veneer (WOOD).

**Figure 8.**
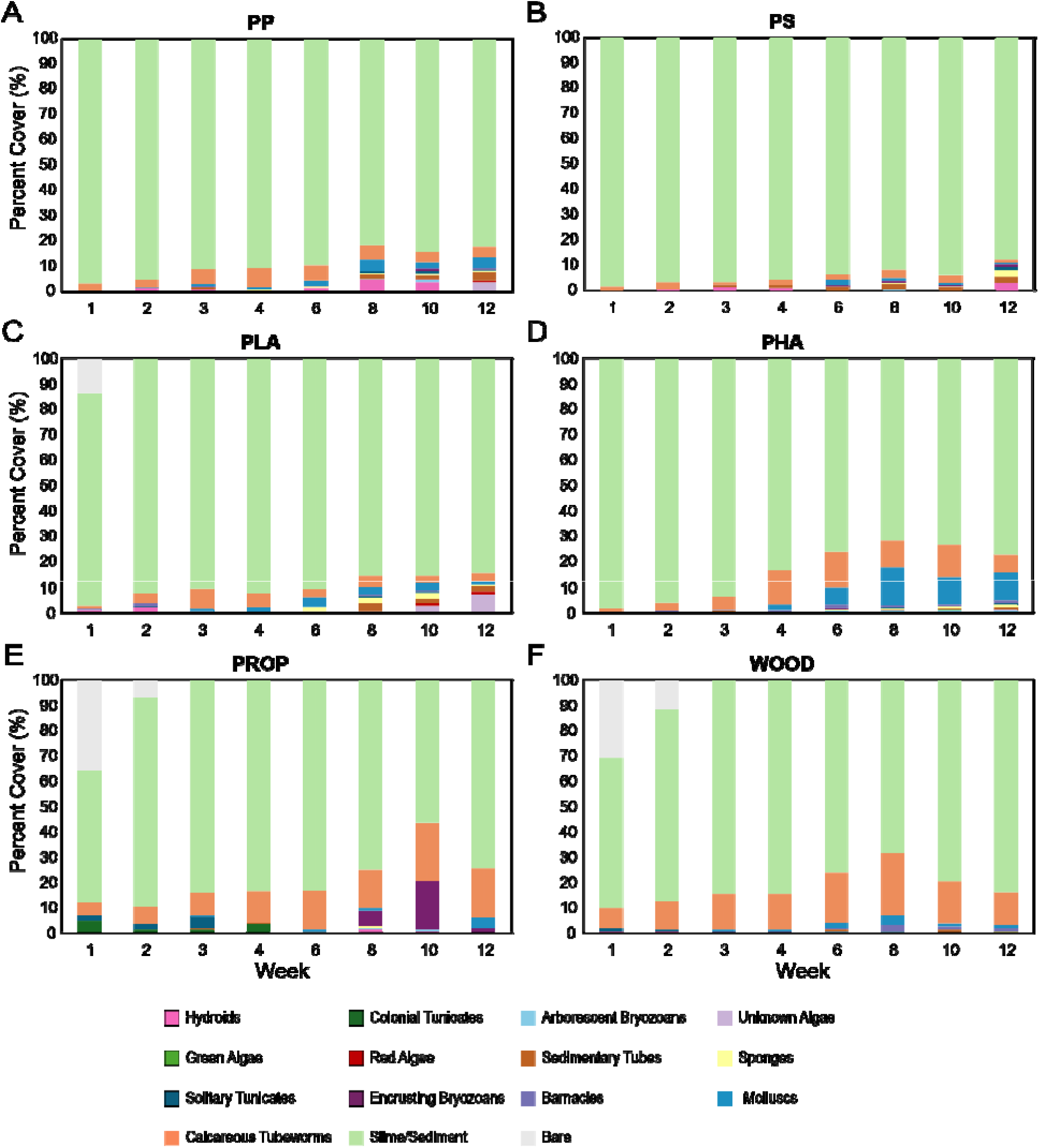
Relative Abundance of Macrofouling Communities. Stacked bar plots depicting the relative abundance of macrofoulers observed on Polypropylene (PP; A), Polystyrene (PS; B), Polylactic Acid (PLA; C), Polyhydroxyalkanoate (PHA; D), mangrove propagules from *Rhizophora mangle* (PROP; E), and wood maple veneer (WOOD; F) at multiple time points throughout the twelve weeks of the study. Colors represent the different macrofoulers present.

The abundance of foulers on all surfaces generally increased over time, though week ten displayed a decline on PHA, wood, and the propagule (Figure 8). Bare surfaces were observed only on the PLA, wood, and propagule and occurred only during the initial weeks of the trial (Figure 8). The wood, mangrove propagule, and PHA recruited more calcareous tubeworms than the other surfaces, despite calcareous tubeworms being a dominant organism on all substrates. *Hydroides elegans* were the most prevalent type of calcareous tubeworm observed. PHA recruited more molluscs than any other substrate (Figure 8).

## 4. Discussion

### 4.1 Biodegradable Plastics Show Variable Rates of Degradation

In the research reported here, PLA and PHA displayed differential degradation patterns under the same marine conditions. PLA showed signs of degradation as early as 22 weeks of immersion in the ocean. This finding contrasts with that of Odobel et al. (2021), who found that PLA did not degrade after seven months of immersion in tanks with flowing seawater. The timescale and rate of plastic degradation depend on the material’s chemical and physical properties, environmental conditions, and other potential factors (Tokiwa et al., 2009). Such discrepancies suggest that biodegradable materials often require specific environmental conditions for optimal degradation. There is also recent evidence that several bacteria can aid in the fragmentation of plastic polymers in the marine environment (Lv et al., 2024). Jiao et al. (2024) noted that multiple marine bacteria were associated with PLA degradation. In our study, the exposure conditions may have been more favorable for PLA biodegradation than for PHA.

In contrast to PLA, PHA required more force to penetrate it after 22 weeks in the ocean. Although this increase in force was not significant, it still may suggest photodegradation by UV radiation. Photodegradation begins with the production of free radicals that can impact plastic in two ways: by causing chain scission to shorten polymer chains or by altering cross-linkages between them. These changes can make the material less flexible, potentially embrittling PHA, and causing it to be stiffer (Ibrahim, 2024). Enzymes capable of degrading PHA are commonly found in the ocean, likely because PHA can be naturally produced by marine microorganisms (Hachisuka et al., 2023; Suzuki et al., 2020). A meta-analysis by Dilkes-Hoffman et al. (2019) estimated that a bottle made from PHA would take approximately 1.5-3.5 years to degrade in the marine environment. Extending the experimental window of time that the substrates are immersed in the ocean might yield a more complete synthesis of the PHA degradation cycle. PHA may recruit more degradation-associated bacteria, given the prevalence of PHA-degrading bacteria in the ocean (Hachisuka et al., 2023; Suzuki et al., 2020).

### 4.2 Microbiome Analyses and Metagenomics

The alpha diversity trends recorded here suggest that biodegradable plastics and biological substrates recruit more complex microbial communities than non-biodegradable plastics and support the role of substrate type in influencing both the composition and diversity of biofilms that form on rafting communities. One potential source of these differences in microbial biofilm composition across substrates may be chemical leachates from the plastics themselves. Many commercial plastics release potentially toxic chemicals into the water during the early stages of immersion (Focardi et al., 2022; Peng et al., 2024; Zimmermann et al., 2021). These leachates can affect local conditions by providing additional organic substrates, trace compounds, and chemicals that may be toxic or alter the behaviors of certain microbial taxa (Focardi et al., 2022; Peng et al., 2024; Zimmermann et al., 2021). Even low concentrations of these compounds can create micro-gradients on or near the substrate surface, thereby shaping early colonization dynamics and selecting microbes with specific metabolic or tolerance traits (Antunes et al., 2019). As a result, the observed community patterns may reflect not only the physical properties of the substrates but also the chemical environments generated by their leachates or by leachates from adjacent debris in the water column.

Additionally, this study revealed that the composition of the biofilms that developed on the two biodegradable plastics, PLA and PHA, recruited different microbial profiles in Pearl Harbor, which is likely due to variation in their polymer chemistry. In a similar study, Marín et al. (2023) investigated PLA and a copolymer within the PHA family, polyhydroxybutyrate-co-hydroxyvalerate (PHBV), and observed distinctly different bacterial compositions in the biofilms that formed on each of the two polymer types. Additionally, biofilms on PHBV produced many of the proteins known to be involved in its biodegradation, whereas biofilms on PLA did not (Marín et al., 2023). PLA and PHA are amongst the most commonly produced biodegradable plastics, and their production is projected to increase. In this study, the biofilms on PHA more closely resembled those on wood and propagules. This finding warrants closer study as it could suggest that marine rafting communities that form on PHA are more consistent with naturally occurring raft communities.

Cyanobacteriota made up the dominant bacterial phylum across all treatments, suggesting an important role for these bacteria in both early- and late-stage biofilm formation at Pearl Harbor. This observation aligns with other studies that have identified Cyanobacteriota as a key bacterial phylum in plastic-associated biofilms in aquatic environments (Amaral-Zettler et al., 2015; Li et al., 2025; Seeley et al., 2020; Zettler et al., 2013). The results reported here found Cyanobacteriota to be dominant, with eleven of the top eighteen identified bacterial genera belonging to the phylum Cyanobacteriota (Figure 5). These include the following bacterial genera: *Prochlorococcus, Fischerella, Leptochromothrix, Cyanothece, Komarekiella, Limnospira, Purpureonostoc, Mojavia, Halarcobacter, Arcobacter, Nostoc, Paracoccus, Selenomonas, Euryhalinema, Phyllonema, and Desmonostoc*. This finding suggests the plastisphere microbial community is shaped by substrate-specific and environmental factors that favored the colonization of Cyanobacteriota over other types on the surfaces investigated in our study. Wallbank et al. (2025) recently reported that plastic polymer type does not significantly drive the associated biofilm community composition, based on studies comparing various plastic types with glass and seawater. However, their findings are not directly comparable to those of our study, because they worked in the temperate region of Auckland with sea temperatures between 15-22°C, while our study took place in Hawai‘i, which is within a tropical region and has average sea temperatures between 24-27°C (Chappell, 2013; Pacific Islands Ocean Observing System (PacIOOS), n.d.).

In contrast to the works mentioned above, the dominance of Cyanobacteriota across all treatments is inconsistent with results in previous studies of marine biofilms sampled from glass slides in Pearl Harbor (Lema et al., 2019; Vijayan & Hadfield, 2020). These researchers found that the most abundant bacterial phylum in both studies was Alphaproteobacteria. Furthermore, Cyanobacteriota were not among the top groups for either study. This disparity between our results and those in earlier investigations may be due to differences in the substrates examined; both Lema et al. (2019) and Vijayan and Hadfield (2020) used glass slides as substrates, whereas we investigated biofilm formation on plastics and biological materials. Another plausible explanation for the differences observed in the microbial communities may be the time of year the biofilms were sampled. Biofilm sample collection took place in October 2023 and January 2024, and the water temperatures in Pearl Harbor vary approximately 4°C between winter and summer months, e.g., 24 - 28°C (Coles, 2006; Ward et al., 2023). The study by Lema et al. (2019) sampled biofilms from Pearl Harbor in July, and the study by Vijayan and Hadfield (2020) sampled them in April, while our study sampled biofilms in October at 27°C and in January at 24°C.

Among the bacterial genera observed, *Alteromonas* and *Paracoccus*, both Pseudomonodotans, stand out for their potential role in plastic degradation. Barbe et al. (2024) showed that *Alteromonas* strains isolated from marine plastic debris incubated in an aquarium with flowing seawater (NW Mediterranean Sea) were present in biofilms on PHBV, a co-polymer of PHA, and aerobically mineralized the material. Other bacteria within the phylum Pseudomonadota have been associated with the ingestion of petroleum-based plastics (Lv et al., 2024). For example, Kim et al. (2020) reported that polystyrene can be degraded by a species of Pseudomonadota isolated from the larvae of a coleopteran. As discoveries of bacteria that can degrade and fragment both bio- and petroleum-based plastics increase, our understanding of the role of bacterial biofilms in the persistence of plastic in our oceans will become clearer.

### 4.3 Colonization by Marine Invertebrates

Non-biodegradable plastics, PP and PS, and the biodegradable plastic, PLA, were observed to have fewer marine invertebrates overall but greater variation in species present than PHA, wood, or the mangrove propagule (Figure 8). This observation suggests that more stable and diverse invertebrate communities can be supported by PHA and biological materials than by non-biodegradable plastics and PLA. The differences observed in the macrofouling communities are likely correlated with the respective microbial communities present on the surface. Marine invertebrate larvae rely primarily on cues produced by the bacteria present within biofilms to settle and metamorphose (Cavalcanti et al., 2020; Dobretsov & Rittschof, 2020; Hadfield & Paul, 2001; Hadfield, 2011; Hadfield et al., 2025). The composition of bacterial biofilms will determine which types of fouling organisms settle on the surfaces.

## 5. Conclusion

The findings reported here demonstrate that biodegradation of plastics in the marine environment is not uniform, but instead depends on polymer type, environmental conditions, surface properties, and the composition of associated microbial biofilms. The contrasting behavior of PLA and PHA in this study highlights that materials labeled as “biodegradable” do not degrade consistently under natural conditions, exposing a key limitation of current standards, which are largely based on controlled laboratory testing. These results underscore the need for field-relevant testing frameworks that better capture the complexity of real-world environments. This is particularly important given that many biodegradable plastics are unlikely to enter specialized recycling streams and instead may be improperly disposed of into natural systems. As such, the effectiveness of biodegradable plastics as a mitigation solution to marine plastic pollution depends not only on their material properties but also on their ability to degrade under environmentally realistic conditions. Materials that support microbial communities capable of efficiently breaking down polymer structures may offer greater potential for rapid conversion into biomass, reducing persistence and the risk of ecological impacts such as transport of invasive species. Moving forward, the design and regulation of biodegradable plastics should prioritize both environmental performance and practical disposal pathways to more effectively reduce plastic pollution and protect marine ecosystems.

## CRediT authorship contribution statement

KMM: Conceptualization, Methodology, Software, Validation, Formal Analysis, Investigation, Resources, Data Curation, Writing – Original Draft, Visualization, Funding Acquisition. BTN: Conceptualization, Methodology, Validation, Resources, Writing - Review & Editing, Supervision. MGH: Conceptualization, Validation, Resources, Writing - Review & Editing, Supervision, Project Administration, Funding Acquisition. MLF: Conceptualization, Software, Validation, Formal Analysis, Data Curation, Writing - Review & Editing, Visualization, Supervision

## Acknowledgements

We would like to thank the University of Hawai‘i at Mānoa Kewalo Marine Lab Faculty, especially Dr. Kiana Frank and Dr. Robert Richmond, for access to their analytical facilities. We also want to thank Meghan Parker and the team at Ho□omalu Ke Kai for their assistance in polymer verification. Lastly, we would like to acknowledge lab members, friends, and family who offered their support and encouragement during this project, specifically Mrs. Amy Knowles and Mr. Andrew Montpetit.

