## Supplemental Materials for "Biodegradable Plastics in the Marine Environment: Comparative Analyses of Bacterial Biofilms, Macroinvertebrate Colonization, and Degradation Patterns"

**Supplementary Table S1. *Macrofoulers Abbreviations.*** Abbreviations and descriptions for classes of fouling organisms commonly observed at the Ford Island Test Site. H/S = Hard or Soft fouling organism

| **Abbreviation** | **H/S** | **Description** |
| --- | --- | --- |
| BAR | H | Barnacles |
| ABRY | H | Arborescent bryozoans |
| EBRY | H | Encrusting bryozoans |
| MOL | H | Bivalve mollusks, mainly oysters |
| CALCTUBE | H | Tube-building polychaete worms, calcareous tubes, mainly *Hydroides elegans* |
| SEDTUBE | S | Tube-building polychaetes and crustaceans, sediment tubes |
| HYD | S | Hydroids |
| UNKA | S | Unknown macroalgae |
| MAR | S | Red macroalgae |
| MAG | S | Green macroalgae |
| MAB | S | Brown macroalgae |
| SPON | S | Sponges |
| CTUN | S | Colonial tunicates |
| STUN | S | Solitary tunicates |
| SLIME |  | Sediment, biological slimes, bare, or unidentifiable |

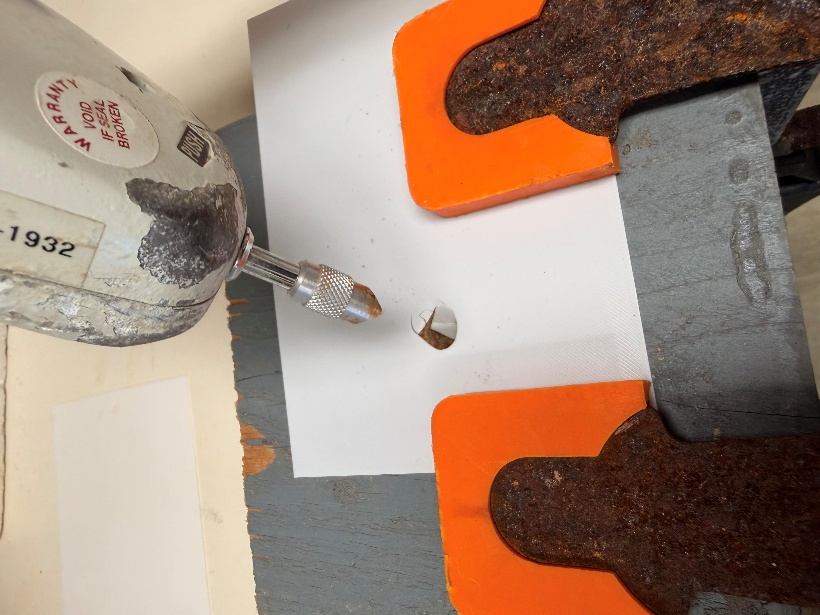

**Supplementary Figure S1. *Force Gauge Measurement Design.*** An image depicting the methodology used for force gauge degradation testing using a chisel tip (0.28 x 8.91 mm) on a mechanical force gauge (SHIMPO).

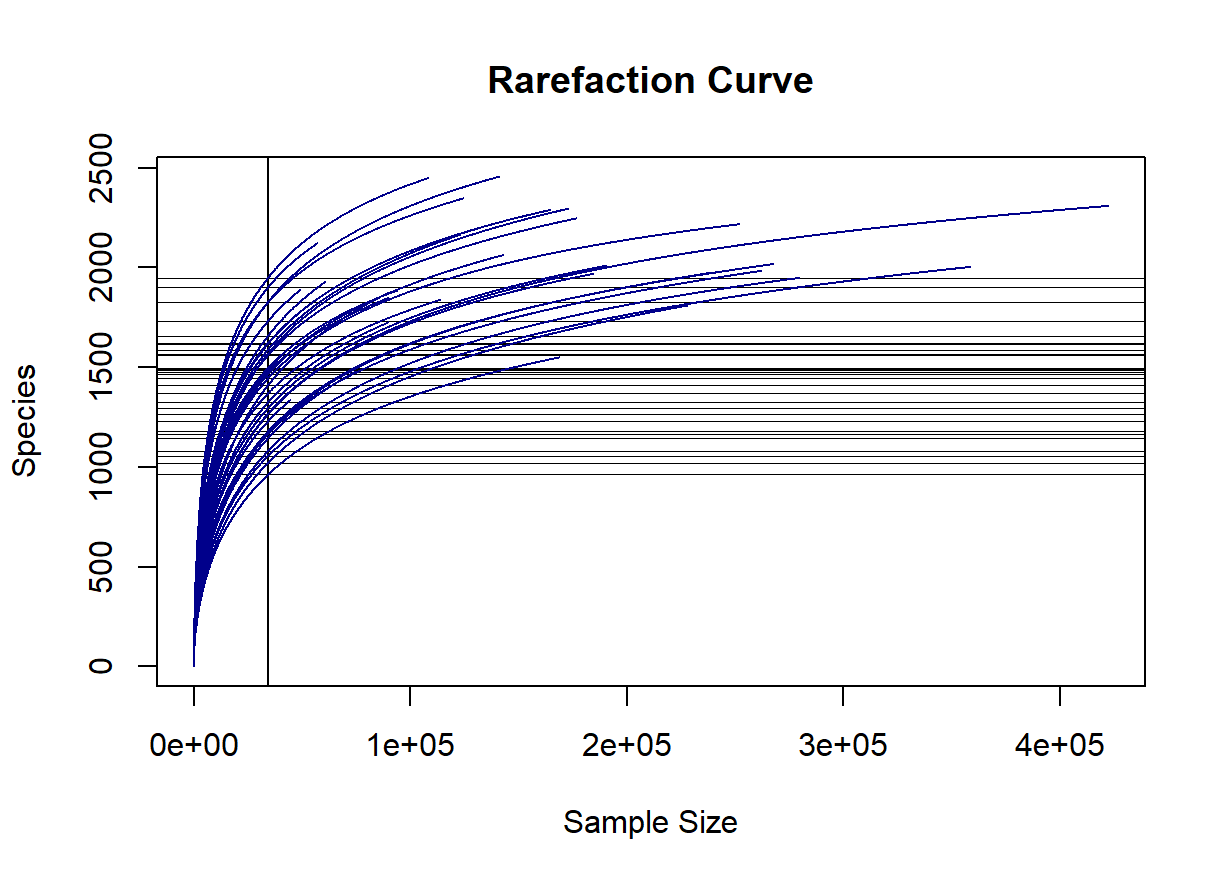

**Supplementary Figure S2. *Rarefaction Curve for Microbial Community Analysis.*** A rarefaction curve conveying the sample size required to be representative of the microbial diversity present. The vertical line is the 34,190-sample effort used for the NMDS and Adonis analyses. The horizontal lines indicate the richness of rarefied species.

**Supplementary Table S2.1 *Results of Force Gauge ANOVA. (Model: force ~ time * substrate)***

|  | **Sum Sq** | **Mean Sq** | **NumDF** | **DenDF** | **F value** | **Pr(>F)** | **P val** |
| --- | --- | --- | --- | --- | --- | --- | --- |
| Time | 1943.03 | 1943.03 | 1 | 28 | 41.2556 | 5.908e-07 | <0.001 |
| Substrate | 669.84 | 111.64 | 6 | 28 | 2.3704 | 0.0560860 | 0.1 |
| Time:Substrate | 1705.15 | 284.19 | 6 | 28 | 6.0341 | 0.0003806 | <0.001 |

**Supplementary Table S2.2 *Results of Pairwise Analysis of Force Gauge Data. (Tukeys)***

| **Substrate** | **Contrast** | **Estimate** | **SE** | **df** | **t.ratio** | **p.value** |
| --- | --- | --- | --- | --- | --- | --- |
| PET | POST - PRE | -6.303 | 5.6 | 28 | -1.125 | 0.2702 |
| PP | POST - PRE | -0.393 | 5.6 | 28 | -0.070 | 0.9445 |
| PLA | POST - PRE | -13.1 | 5.6 | 28 | -2.338 | 0.0268 |
| PHA | POST - PRE | 3.570 | 5.6 | 28 | 0.637 | 0.5292 |
| PROP | POST - PRE | -33.740 | 5.6 | 28 | -6.021 | <0.0001 |
| WOOD - with | POST - PRE | -17.863 | 5.6 | 28 | -3.188 | 0.0035 |
| WOOD - against | POST - PRE | -27.393 | 5.6 | 28 | -4.889 | <0.0001 |

**Supplementary Table S3.1 *Results of* *ANOVA of Shannon Diversity Indices. (Shannon ~ Substrate * Time)***

|  | **Df** | **Sum Sq** | **Mean Sq** | **F value** | **Pr(>F)** |
| --- | --- | --- | --- | --- | --- |
| **Substrate** | 5 | 4.067 | 0.813 | 3.352 | 0.022150 |
| **Time** | 1 | 4.301 | 4.301 | 17.719 | 0.000394 |
| **Substrate:Time** | 5 | 1.108 | 0.222 | 0.913 | 0.491672 |
| **Residuals** | 21 | 5.097 | 0.243 |  |  |

**Supplementary Table S3.2 *Results of* *Pairwise Comparisons of Shannon Dive*rsity Indices. *(Tukeys)***

| **Comparison** | **diff** | **lwr** | **upr** | **p adj** |
| --- | --- | --- | --- | --- |
| PLA-PHA | -0.4160927 | -1.34938204 | 0.51719659 | 0.7295946 |
| PP-PHA | -0.8566347 | -1.78992397 | 0.07665467 | 0.0841304 |
| PROP-PHA | -0.1520578 | -1.04191424 | 0.73779866 | 0.9940336 |
| PS-PHA | -0.6684816 | -1.60177089 | 0.26480775 | 0.2614930 |
| WOOD-PHA | 0.1283027 | -0.76155374 | 1.01815915 | 0.9973048 |
| PP-PLA | -0.4405419 | -1.41533082 | 0.53424698 | 0.7186553 |
| PROP-PLA | 0.2640349 | -0.66925438 | 1.19732426 | 0.9459652 |
| PS-PLA | -0.2523888 | -1.22717775 | 0.72240005 | 0.9624517 |
| WOOD-PLA | 0.5443954 | -0.38889389 | 1.47768475 | 0.4721530 |
| PROP-PP | 0.7045769 | -0.22871245 | 1.63786618 | 0.2144908 |
| PS-PP | 0.1881531 | -0.78663582 | 1.16294198 | 0.9895820 |
| WOOD-PP | 0.9849374 | 0.05164804 | 1.91822667 | 0.0347376 |
| PS-PROP | -0.5164238 | -1.44971310 | 0.41686553 | 0.5277340 |
| WOOD-PROP | 0.2803605 | -0.60949596 | 1.17021694 | 0.9174519 |
| WOOD-PS | 0.7967843 | -0.13650504 | 1.73007359 | 0.1237303 |

**Supplementary Table S4*. PERMANOVA Analysis Comparing Substrate Type (Biodegradable, Biological, Non-Biodegradable)***

|  | **Df** | **Sum of Sqs** | **R^2^** | **F** | **Pr(>F)** |
| --- | --- | --- | --- | --- | --- |
| Model | 5 | 1.3282 | 0.4142 | 3.8182 | 0.001 |
| Residual | 27 | 1.8784 | 0.5858 |  |  |
| Total | 32 | 3.2066 | 1.0000 |  |  |
